# Estrogen and Soy Isoflavonoids Decrease Sensitivity of Medulloblastoma and Central Nervous System Primitive Neuroectodermal Tumor Cells to Chemotherapeutic Cytotoxicity

**DOI:** 10.1101/100107

**Authors:** Scott M. Belcher, Caleb C. Burton, Clifford J. Cookman, Michelle Kirby, Gabriel L. Miranda, Fatima Saeed, Kathleen E. Wray

## Abstract

**Background:** Our previous studies demonstrated that growth and migration of medulloblastoma (MB), the most common malignant brain tumor in children, are stimulated by 17β-estradiol. The growth stimulating effects of estrogens are mediated through ERβ and insulin-like growth factor 1 signaling to inhibit caspase 3 activity and reduce tumor cell apoptosis. The objective of this study was to determine whether estrogens decreased sensitivity of MB cells to cytotoxic actions of chemotherapeutic drugs.

**Methods:** Using *in vitro* cell viability and clonogenic survival assays, concentration response analysis was used to determine whether the cytoprotective effects of estradiol protected human D283 Med MB cells from the cytotoxic actions of the MB chemotherapeutic drugs cisplatin, vincristine, or lomustine. Additional experiments were done to determine whether the ER antagonist fulvestrant or the selective ER modulator tamoxifen blocked the cytoprotective actions of estradiol. ER-selective agonists and antagonists were used to define receptor specificity, and the impacts of the soy-derived phytoestrogens genistein, daidzein, and s-equol on chemosensitivity were evaluated.

**Results:** In D283 Med cells the presence of 10 nM estradiol increased the IC_50_ for cisplatin-induced inhibition of viability 2-fold from ~5 μM to >10 μM. In clonogenic survival assays estradiol decreased the chemosensitivity of D283 Med exposed to cisplatin, lomustine and vincristine. The ERβ selective agonist DPN and low physiological concentrations of the soy-derived phytoestrogens genistein, daidzein, and s-equol also decreased sensitivity of D283 Med cells to cisplatin. The protective effects of estradiol were blocked by the antiestrogens 4-hydroxytamoxifen, fulvestrant (ICI 182,780) and the ERβ selective antagonist PPHTP. Whereas estradiol also decreased chemosensitivity of PFSK1 cells, estradiol increased sensitivity of Daoy cell to cisplatin, suggesting that ERβ mediated effects may vary in different subtypes of MB.

**Conclusions:** These findings demonstrate that E2 and environmental estrogens decrease sensitivity of MB to cytotoxic chemotherapeutics, and that ERβ selective and non-selective inhibition of estrogen receptor activity blocks these cytoprotective actions. These findings support the therapeutic potential of antiestrogen adjuvant therapies for MB, and findings that soy phytoestrogens also decrease sensitivity of MB cells to cytotoxic chemotherapeutics suggest that decreased exposure to environmental estrogens may benefit MB patient responses to chemotherapy.

## Background

Medulloblastoma (MB) arise from neural precursors of the cerebellum and brainstem and are associated with the 4^th^ ventricle. They are the most common central nervous system (CNS) malignancy in childhood with a peak incidence around 5 years of age [1–4]. These primitive neuroectodermal tumors (PNETs) while rare, with an overall incidence rate of 1.5 per million population, are more common in children 1-9 years of age (affecting 9.6 per million children) compared to adults 19 years of age and older, who have an incidence rate of 0.6 per million [5]. Less commonly, PNETs may develop in the cerebral hemisphere, these tumors are referred to CNS-PNETs. While sharing histological similarities, CNS-PNET tumors are genetically distinct from MB and have an overall incidence rate of 0.62 per million [5–8]. Histopathology grading has classically been used to separate MB into subgroups which differ with regard to biomarker profile and prognosis [9]. These subgroups include classic MB, desmoplastic MB, MB with excessive nodularity (MBEN), large cell MB and anaplastic MB [9–11]. Due to cellular and molecular heterogeneity across histological subgroups, and even within a singular tumor, a newer approach to MB grading has emerged that relies on comparative genome, transcriptome, and epigenetic analysis which may allow improved risk stratification and individualized targeted treatments [12]. By consensus four molecular subgroups are now recognized, they include wingless (WNT), sonic hedgehog (SHH), group 3 and group 4 [12–15]. Each subgroup has a characteristic genetic profile and gene expression patterns that appear to drive tumor progression, predict therapeutic responsiveness and prognosis. Further refinement of these molecular analyses has also found that pediatric and adult MB, are both histologically and genetically different diseases with characteristic differences in mutation accumulation, chromosomal deletion and amplification and distinctive prognosis and survival rates [16–18].

Advancements in multimodal MB therapy utilizing maximal tumor resection, followed by radiation, and chemotherapy have greatly improved the chances of patient survival with 5 year overall survival rates for MB reaching between 60-80% depending on specific tumor grade or molecular subtype; the survival rate for CNS-PNET patients is approximately 50% [18]. Conventional standard of care for MB most often involves combined radiation and polychemotherapy that results in improved outcomes compared to treatments limited to only tumor excision, radiation therapy or single agent chemotherapy [8, 19, 20]. Cytotoxic chemotherapy treatments for standard risk MB include a combination of cisplatin, cyclophosphamide, lomustine, and vincristine [21]. These agents vary in their mechanism of actions with cisplatin causing apoptosis due to DNA cross-linking, cyclophosphamide and lomustine are DNA alkylating agents, and the vinca alkaloid vincristine inhibits cell division by binding tubulin to inhibit microtubule formation [21]. Despite the success of these combined treatments, greater than 70% of MB survivors experience life-long neurological disabilities that include cognitive, motor, and/or vision impairments, as well as psychosocial dysfunction. Additionally, more than half of survivors also have severe endocrine impairments, which further contribute to a greatly diminished quality of life for MB survivors [22–24]. Thus, there is continued need to refine existing therapy and develop new adjuvant therapies that further improve MB and CNS-PNET cure rates, while reducing the life-long adverse effects of both the disease and its treatment [25].

Previous study has demonstrated that growth and migration of MB and CNS-PNET cells and tumors are responsive to estrogen (17β-estradiol; E2) and other estrogenic compounds [26–30]. In some human MB cell lines and in genetic mouse models of MB, the growth stimulating effects of estrogen are mediated through ERβ regulation of prosurvival insulin-like growth factor 1 (IGF1) signaling pathways that acts to reduce tumor cell apoptosis [26–28]. Results from additional studies have also demonstrated that therapeutic doses of the ER antagonist fulvestrant inhibited MB cell growth and migration in cultured human MB cell lines, blocked the growth of MB tumors in nude mice, and decreased tumor growth and progression in mouse genetic models of MB that most closely resemble the SHH subtype of MB [26–28]. These findings support the potential efficacy of anti-estrogen treatments, or other interventions to decrease ER activity, as potentially beneficial adjuvants for MB management. The role of ERβ in MB pathology however, is controversial in part because the role of ERβ in cancer progression is in general poorly understood, with both tumor suppressor and tumor promoting effects of ERβ having been reported in different ER expressing tumors [31, 32]. Results from different mouse models of MB have also found that estrogen and ERβ activity can decrease MB tumor incidence [33], and *in vitro* studies have suggested that in the Daoy MB cell line, inhibition of ERβ activity decreases sensitivity to cisplatin by enhancing Rad51 mediated DNA repair mechanisms [34]. To investigate the role of estrogens in MB in more detail, we have used cell viability and clonogenic survival assays to determine whether the cytoprotective effects of E2 protected human D283 Med MB cells from the cytotoxic actions of the MB chemotherapeutic drugs cisplatin, vincristine, or lomustine. The effects of E2 on cisplatin chemosensitivity were also determined in the MB cell line Daoy and a CNS-PNET cell line PFSK1. Additional experiments were done to determine whether the ER antagonist fulvestrant or the selective ER antagonist tamoxifen blocked the cytoprotective actions of 17β-estradiol, and whether other ER selective agonists, and low concentrations of the soy-derived phytoestrogens genistein, daidzein, and s-equol were able to impact D283 Med chemosensitivity.

## Materials and Methods

### Steroids and Pharmacological Agents

Dimethylsulfoxide (DMSO), 3-(4,5-Dimethyl-2-thiazolyl)-2,5-diphenyl-2H-tetrazolium bromide (MTT), 4-hydroxytamoxifen (4-OHT), 4’7-dihydroxyisoflavonoid (daidzein), 4’,5,7-trihydroxyisoflavonoid (genistein) and 17β-estradiol (E2) were from Sigma-Aldrich (St. Louis, MO). 4′,7-dihydroxyisoflavan (s-equol) was from Cayman Chemical (Ann Arbor, MI). Fulvestrant (ICI 182, 780), 4,4′,4″-(4-propyl-[1*H*]-pyrazole-1,3,5-triyl)*tris*phenol (PPT), 2,3-*bis*(4-Hydroxyphenyl)-propionitrile (DPN) and 4-[2-Phenyl-5,7-*bis*(trifluoromethyl)pyrazolo [1,5-*a*]pyrimidin-3-yl]phenol (PHTPP) were from Tocris Bioscience (R&D Systems, Inc., Minneapolis, MN). Cisplatin, vincristine, and lomustine were from Selleck Chemical (Houston, TX). Cisplatin was usually prepared as a 1 mg/mL (3.3 mmole/L) stock solution in PBS, in assays involving hydrophobic ligands in DMSO results were normalized using a standard curve comparing D283 Med cytotoxicity in the presence or absence of DMSO [35].

### Cell Culture Conditions

All cell lines were acquired from the American Type Culture Collection, cryopreserved and expanded for analysis. The unique growth and morphological characteristics of each cell line was retained throughout the duration of the study. Details of cell culture methods were described previously [26–28]. Briefly, D283 Med and Daoy cells were grown in a humidified incubator at 37°C and 5% CO_2_ atmosphere in growth media containing minimum essential media (MEM) with Earle’s Balanced Salt Solution (EBSS). Growth media for PFSK1 cells was RPMI 1640. Media was supplemented with 10% fetal bovine serum (FBS), 4 mM L-glutamine, 100 U/ml penicillin and 100 μg/ml streptomycin (Thermo Fisher Scientific, Waltham, MA). For general growth and expansion, D283 Med cells were maintained in suspension culture at 0.5 – 1 × 10^6^ cells/ml. Daoy and PFSK1 cells were maintained between 20 and 80% confluence. Growth media was refreshed every 2-3 days with cells split at a ratio of 1:5.

For Daoy growth analysis cells from subconfluent cultures were harvested by dissociation with 0.2 mM EDTA in PBS, resuspended in phenol-free supplemented with in 10% charcoal stripped FBS (CSS) and viable cell numbers determined by direct cell counting of trypan blue-excluding cells with a hematocytometer. Cells were seeded in triplicate into 60 mm culture dishes (22.06 cm^2^). Optimization experiments with cells plated at an initial density of 1,000, 3,000, 10,000, or 20,000 cells per dish in 10% FBS, 10% CSS plus or minus various concentrations of E2 indicated that 3000 cells per well allowed optimal viability analysis at all time points [28]. Cultures were untreated, or treated with DMSO (0.001%), 10 nM E2, that was serially diluted into fresh DMSO/PBS vehicle to obtain an equal 0.001% final DMSO concentration in all cultures, and 10% FBS served as a positive control. At 24, 48, 72 and 96 hours post-treatment viable cell numbers were determined by direct counting of trypan blue-excluding cells.

### Viability Analysis

Viability was assessed by accumulation of formazan by reduction 3-(4,5-dimethylthiazol-2-yl)-2,5-diphenyltetrazolium bromide (MTT) or 3-(4,5-dimethylthiazol-2-yl)-5-(3-carboxymethoxyphenyl)-2-(4-sulfophenyl)-2H-tetrazolium (MTS) in the presence of phenazine methosulfate (CellTiter 96^®^ AQueous One Solution Cell Proliferation Assay; Promega) as previously described [36]. To avoid any potential MTT/MTS reduction assay bias, effects of E2 on D283 Med cell viability were confirmed with separate experiments using an Alamar Blue (resazurine) fluorescent dye assay at excitation wavelength of 535 nm (20 nm bandwidth) and an emission wavelength of 590 nm (35 nm bandwidth) [37]. Comparable results were observed for all assays. Regardless of specific assay, growing cells were harvested, counted and resuspended at a desired density in 10% CSS supplemented MEM/EBSS with 10 nM E2, or desired final concentration of fulvestrant or the vehicle control (0.01% DMSO) prior to cisplatin exposure. Cultures were incubated at 37°C in 5% CO_2_ overnight (18-24 hours) at which time cells were exposed to the desired final concentration of cisplatin and incubated an additional 48 hours prior to viability analysis. For each bioassay D283 Med cells were seeded in 96 well plates at 1 × 10^5^ cells/mL (1 - 2 × 10^4^ cells per well) based on results of preliminary experiments to optimize each assay.

### Clonogenic Assay

Clonogenic/colony forming assays were adapted from published protocols [38, 39]. Cell were seeded at 1000 cells per well in 6-well tissue culture plates in 2 mL of 10% CSS supplemented phenol red free MEM/EBSS. For D283 Med cells poly-L-lysine coated culture plates were used allowing adherent growth. Exposure to ER ligands were started 24 hours prior to chemotherapeutic exposure and continued during and following chemotherapeutic agent exposure. For chemotherapeutic drug exposures, 0.5 mL of a 5x stock prepared in cell culture media was added to each well. After 6 hours (cisplatin) or 24 hours (vincristine and lomustine) of exposure, media was aspirated, cells were washed 2 times with chemotherapeutic compound-free media, and then cultured in 2.5 mL of growth media at 37°C in 5% CO_2_ until visible colonies containing >50 cells were observed. Preliminary range finding concentration response analysis was performed with each chemotherapeutic agent for each cell line, at the concentrations used between 5-40 clones per plate were typically observe for vehicle cultures. Incubation times were typically between 2-3 weeks with growth media refreshed every 2-3 days. Colonies were fixed and stained with 1% methylene blue in 50% ethanol (Fisher Scientific, Pittsburg, PA) or 0.1% coomassie brilliant blue in methanol (Bio-Rad, Hercules, CA). Digital images were captured and colonies were counted using an Alpha Innotech FluorChem FC2 imager (ProteinSimple, Santa Clara, CA) and Adobe Photoshop (Adobe Systems Inc., San Jose, CA).

### Caspase Activity

All methods were done as previously described with D283 Med cells seeded into 96 well plates at a density of 1 × 10^6^ cells/ml in phenol red free MEM/EBSS lacking L-glutamine, 10% CSS and supplemented with increasing concentrations of daidzein, genistein or s-equol [27]. Cells exposed to 10 nM E2 or DMSO vehicle served as controls. Cells were lysed 48 hours after seeding with 20 mM Tris-HCl (pH 7.5) with 150 mM NaCl, 1 mM EDTA, 1 mM EGTA, and 1% Triton. Cell lysates were assayed for protein concentration using the BioRad Dc protein assay (Bio-Rad, Hercules, CA). Caspase activity (pmol of pNA hour^−1^ mg protein^−1^) from 10 μg of lysate was determined by comparing the amount p-nitroaniline (pNA) liberated from Ac-DEVD-pNA (Enzo Life Sciences, Farmingdale, NY) with a standard curve derived from known concentrations of pNA. Normalized caspase activity for each phytoestrogen are reported as a percentage of the maximal inhibitory effect of 10 nM E2.

### Data and Statistical Analysis

All experiments were repeated a minimum of 3 times. Statistical analysis was conducted using one way ANOVA or two-way ANOVA with Holm-Sidak’s multiple comparisons test. A minimal level of statistical significance for differences between groups was p < 0.05 and unless otherwise noted is indicated by *. Concentration response curves and IC_50_ estimates were generated using a normalized variable slope Hill model. Analysis was performed using GraphPad Prism v6 software (GraphPad Software, Inc., La Jolla, CA).

## Results

Compared to vehicle treated D283 Med cells, the cytotoxic effects of cisplatin were decreased in the presence of 10 nM E2 (Fig. 1A-B). When analyzed by an MTS reduction assay the presence of E2 increased the observed IC_50_ of cisplatin from 5.6 μM (95% CI 4.7 - 6.9) to 14.7 μM (95% CI 10.5 − 20.5; Fig. 1A). Two-way ANOVA revealed a significant effect of cisplatin concentrations [F (5, 36) = 49.65, p < .0001], a significant effect of 10 nM E2 [F (1, 36) = 10.07, p < .0031], and a significant interaction between cisplatin concentration and E2 exposure [F (5, 36) = 5.873, p= 0.0005). Shown in Fig. 1B are results of independent experiments using the resazurin reduction bioassay as an indicator of D283 Med viability where the IC_50_ for cisplatin cytotoxicity in control cultures lacking E2 was calculated as 4.8 μM (95% CI 4.1 - 5.7). Revealing that the observed effects were not an assay specific effect, the presence of 10 nM E2 similarly increased the calculated IC50 for cisplatin cytotoxicity to 9.4 μM (95% CI 7.7 - 11.5). The cytoprotective effects of E2 were blocked by the ER antagonist fulvestrant (ICI 182,780). In the presence of both E2 and fulvestrant, the calculated cisplatin IC_50_ was 4.2 μM (95% CI 3.2 – 5.39) which was indistinguishable from control (Fig. 1B).

**Fig. 1.**
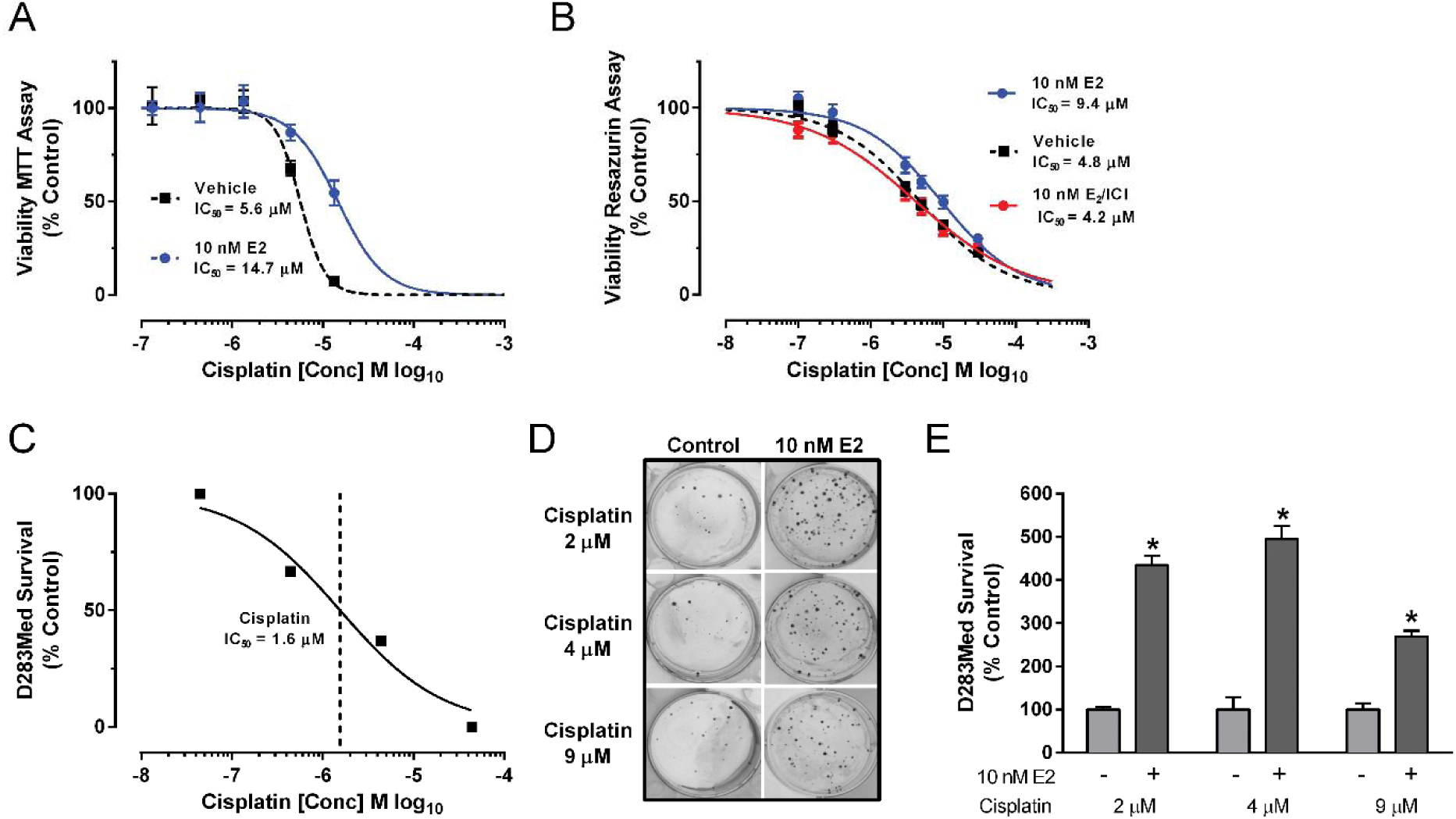
The cytoprotective effect of 10 nM E2 on cisplatin cytoxicity in D283 Med cells. (A) Concentration response analysis of D283 Med viability following exposure to increasing concentrations of cisplatin with and without 10 nM E2 using the MTS-reduction assay. All data is expressed as a percentage (± SEM; n = 4 per dose group) relative to vehicle treated control cultures. Concentration response curves and indicated IC_50_ values for cisplatin inhibition of viability were calculated using a normalized variable slope Hill model. (B) Concentration response analysis of the cytotoxic effects of cisplatin on D283 Med cells in the presence of 10 nM E2 plus or minus 10 nM fulvestrant (ICI 182,780) using the resazurine fluorescent dye assay (for vehicle and E2 groups n = 28 replicates; E2/ICI n = 20 replicates from 3 separate experiments). (C) Initial range finding concentration response analysis of the cytotoxic effects of cisplatin in D283 Med using a colony forming (clonogenic) assay of cell survival defined an IC_50_ of 1.6 μM. (D) Representative images of plates stained with 0.1% coomassie brilliant blue in methanol to visualize colonies formed from cultures of 1000 D283 Med cells in the presence of 10 nM E2 or vehicle control that were treated with 2, 4 or 9 μM cisplatin. (E) Quantification of surviving colony numbers from clonogenic assays of D283 Med cells exposed to 2, 4, or 9 μM cisplatin with or without 10 nM E2 (n = 4 for each group). All results are expressed as mean ± SEM. Significant differences from the control group were determined by two-way ANOVA followed by post-test analysis; * p ≤ .05.

Following the initial characterization studies of the effects of E2 on D283 Med cells in viability assays, a clonogenic colony forming assay [38, 39] in which cytotoxic treatments reproducibly caused 99-99.5% loss of viability was used to better determine the effects of estrogens effects on the cytotoxicity of cisplatin. Based on preliminary concentration response analysis (Fig. 1C), the effect of 10 nM E2 on chemosensitivity of D283 Med cells to increasing cisplatin concentrations (2, 4, or 9 μM) was characterized (Fig. 1D-E). At each cisplatin concentration E2 was significantly cytoprotective [F (1, 18) = 311.6, p < .0001; p < .0001 for each cisplatin concentration]. To determine whether the observed protective effect of E2 in D283 Med cells was independent of the cytotoxic mechanism of action, additional experiments were performed to test the impact of E2 on lomustine and vincristine cytotoxicity (Fig. 2). Initial range-finding concertation response analysis in the D283 Med clonogenic assay indicated an IC_50_ for lomustine of 12.1 μM (95% CI 11.7 – 12.6) (Fig. 2A) and 1.5 nM (95% CI 0.74 – 3.1) for vincristine (Fig. 2B). The presence of E2 significantly protected D283 Med cells from the cytotoxic effect of lomustine [F (1, 30) = 74.64, p < .0001] at each concentration tested (Fig. 2C; 10 μM p = .0036, and p < .0001 for 20 and 40 μM). For vincristine 10 nM E2 also significantly [F (1, 18) = 196.2, p < .0001] decreased cytotoxicity at each concentration (Fig. 2D; 5 and 10 nM, p < .0001 and p = .0003 for 20 nM).

**Fig. 2.**
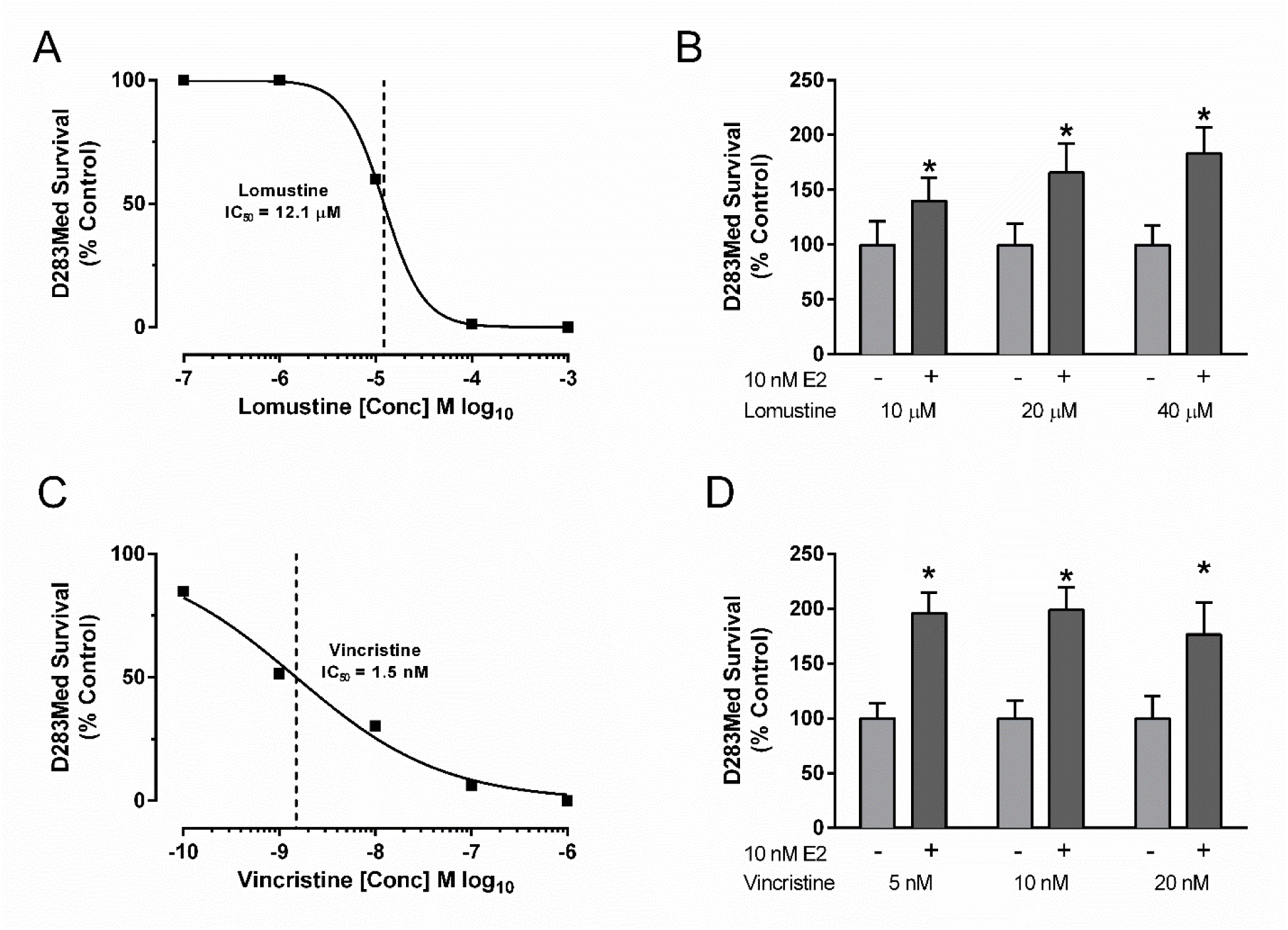
The effects of E2 exposure on lomustine and vincristine cytotoxicity in D283 Med cells. Initial range finding concentration response analysis of the cytotoxic effects of (A) lomustine (IC_50_ = 12 μM) and (B) vincristine (IC_50_ = 1.5 nM) on D283 Med in a clonogenic assay of cell survival. (C) Quantification of surviving colony numbers from clonogenic assays of D283 Med cells exposed to 10, 20, or 40 μM lomustine with or without 10 nM E2 (n = 6 for each group). (D) Quantification of colony number from clonogenic assays of D283 Med cells exposed to 5, 10, or 20 nM vincristine with or without 10 nM E2 (n = 8 for each group). All results are expressed as mean ± SEM. Significant differences from the control group were determined by two-way ANOVA followed by Holm-Sidak’s post-test analysis and is indicated above the error bars; * p ≤ .05.

Compared to vehicle treated D283 Med cells exposed to 4 μM cisplatin, E2 (p < .0001) and the ERβ selective agonist DPN (p < .0001) each increased numbers of surviving colonies compared to cisplatin alone control cultures (Fig. 3A-B). The cytotoxic effect of 4 μM cisplatin was not significantly changed by the ERα selective agonist PPT (p > .9999). At a final concertation of 10 nM, the soy isoflavonoids genistein (p = .0239), daidzein (p < .0001), or the bacterial metabolite of daidzein, (s)-equol (p < .0001) each significantly protected D283 Med cells from the cytotoxic effect of cisplatin (Fig. 3B). The relative magnitude of the protective effects for each of the compounds is consistent with their selectivity and potency at ERβ [40]. Increasing concentrations of each phytoestrogen significantly [F (2, 256) = 4.85, p < .0086] and dose-dependently decreased caspase 3 activity in D283 Med cells. The decrease in caspase activity mirrored the cytoprotective effects seen in the clonogenic assay (Fig. 3B). The differences in the suppression of caspase activity compared to control reached a significant difference in the 10 nM (10^−8^M) groups for s-equol and daidzein and for genistein at 100 nM (Fig. 3C).

**Fig. 3.**
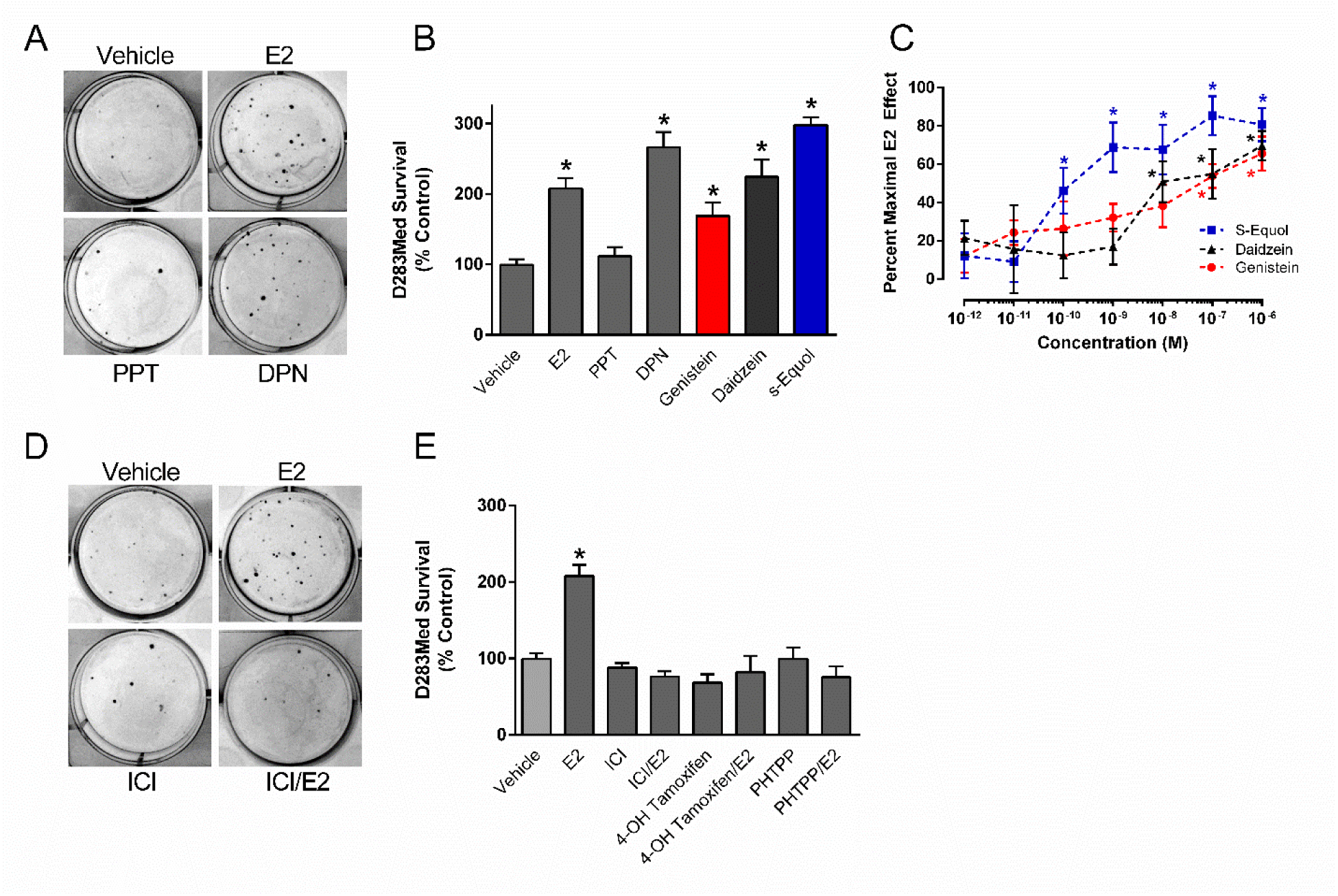
The effects of selective and nonselective ER ligands and soy-derived isoflavonoids on cisplatin cytotoxicity in D283 Med cells. (A) Representative images of surviving coomassie blue stained colonies from D283 Med cultures cotreated with 4 μM cisplatin and either vehicle, 10 nM E2, 10 nM PPT or 10 nM DPN. (B) Quantification of colony number from clonogenic assays of D283 Med cells cotreated with 4 μM cisplatin and vehicle, or 10 nM of E2 (n =18) , PPT (n = 7), DPN (n = 8), or 10 nM of genistein, daidzein, or the daidzein metabolite s-equol (n = 6 for each isoflavonoid group). (C) Concentration response analysis of estrogenic inhibition of caspase 3 activity by increasing concentrations of s-equol, daidzein, or genistein in D283 Med cultures. Control cultures exposed to 0.01% DMSO vehicle or 10 nM E2 were treated and analyzed in parallel. Caspase activity was quantified following a 48 hour incubation period. Results were normalized to the relative caspase 3 activity of the vehicle control and expressed as mean percent of the effects for 10 nM E2. The number of samples in each group was n = 10-12. (D) Representative images of surviving coomassie blue stained colonies from D283 Med cultures cotreated with 4 μM cisplatin and either vehicle, 10 nM E2, 10 nM fulvestrant (ICI), or 10 nM E2 and 10 nM fulvestrant (ICI/E2). (E) Quantification of colony number from clonogenic assays of D283 Med cells cotreated with 4 μM cisplatin and either vehicle (n = 12), 10 nM E2 (n = 12), 10 nM fulvestrant (ICI; n = 8), 10 nM E2 and 10 nM fulvestrant (ICI/E2; n = 7), 1 μM 4-OH tamoxifen with and without 10 nM E2 (n = 8), or 5 μM PHTPP with and without E2 (n = 8). All results are expressed as mean ± SEM. Significant differences from the control group was determined by one-way ANOVA followed by Holm-Sidak’s multiple comparisons tests which is indicated above the error bars: * p ≤ .05.

The cytoprotective effects of E2 in D283 Med cells exposed to cisplatin (p = .0001) were eliminated by the non-selective ER antagonist fulvestrant (10 nM; ICI; E2 vs. E2/fulvestrant p = < .0001; Fig. 3D-E), the selective estrogen receptor modulator 4-OH tamoxifen (1 μM; E2 vs E2/tamoxifen p = < .0001) or the ERβ selective antagonist PHTPP (5 μM; E2 vs E2/PHTPP p = < .0001; Fig. 3E). In CNS-PNET derived PFSK1 cells, 10 nM E2 also resulted in increased survival [F (1, 34) = 62.30, p < .0001], with a clear decrease in sensitivity to the cytotoxic effects observed for all three cisplatin concentrations tested (Fig. 4A). Fulvestrant (10 nM) also blocked the cytoprotective effects of 10 nM E2 (p = .0213; Fig. 4B). In contrast to both D283 Med and PFSK1 cells, the cytotoxic effects of cisplatin were increased in Daoy cells by the presence of 10 nM E2 where a significant decrease [F (1, 18) = 62.75, p < .0001] in surviving colony formation was observed in the estrogen treated cultures (Fig. 4C). The increased sensitivity of Daoy to cisplatin in the presence of E2 (p = .0194) was also eliminated by fulvestrant (p = .0012; Fig. 4D) and 10 nM E2 did not stimulate growth of Daoy cells (Fig. 4E).

**Fig. 4.**
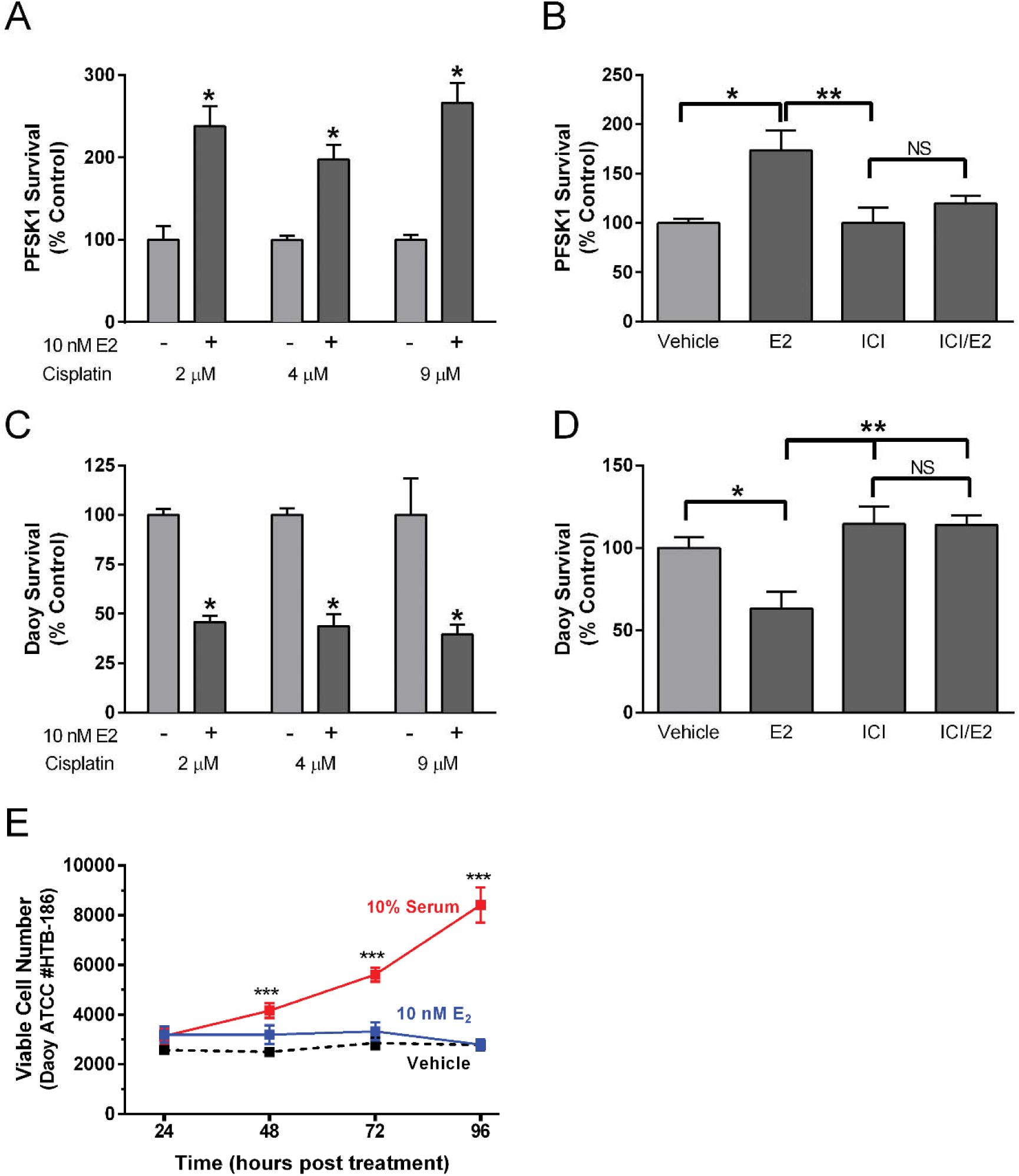
The effects of E2 exposure on cisplatin cytotoxicity in PFSK1 CNS-PNET cells and Daoy cells. (A) Quantification of surviving colony numbers from clonogenic assays of PFSK1 cells exposed to 2, 4 or 9 μM cisplatin with or without 10 nM E2 (n = 8 for each cisplatin treatment group except 4 μM where n = 4). (B) Quantification of colony number from clonogenic assays of PFSK1 cells cotreated with 4 μM cisplatin and either vehicle, 10 nM E2, 10 nM fulvestrant (ICI), 10 nM E2 and 10 nM fulvestrant (ICI/E2), n = 4 for each group. (C) Quantification of surviving colony numbers from clonogenic assays of Daoy cells exposed to 2, 4 or 9 μM cisplatin with or without 10 nM E2. For each group n = 4. (D) Quantification of colony number from clonogenic assays of Daoy cells cotreated with 4 μM cisplatin and either vehicle, 10 nM E2, 10 nM fulvestrant (ICI), 10 nM E2 and 10 nM fulvestrant (ICI/E2), n = 8 for each group. (E) Analysis of the effects of 10 nM E2 on viability of Daoy cells. At T_0_ 3000 cells were plated into 60 mm cell culture dishes, in growth media cells containing 10% CSS. At each indicated time point (hours post treatment) cells were harvested and trypan-excluding cells were counted. Vehicle was 0.0001% DMSO and replacement of CSS with 10% FBS served as a positive control. At each time point n =10 for all treatments. Results are expressed as mean ± SEM. Significant differences from vehicle control are indicated above the treatment group error bars with individual comparisons indicated above brackets: * p ≤.05; ** p ≤ 0.01; *** p ≤ .001; NS, not significant.

## Discussion

The use of aggressive multimodal treatments has resulted in increased survival for MB patients, most survivors however suffer from life-long adverse effects that greatly diminish their quality of life [22]. The presented findings demonstrate that E2 can increase the resistance to cytotoxic chemotherapeutics commonly used to treat MB, and that blockade of estrogen receptor activity inhibits this effect. These findings suggest that ER antagonists may be a useful adjuvant approach to current cytotoxic chemotherapy used to treat MB. The cytoprotective effects of estrogens, either endogenous or derived from environmental sources such as diet or estrogenic endocrine disruptors from medical devices [41, 42], if translatable to MB patients, would require more aggressive chemotherapeutic interventions to achieve a cure in patients with increased levels of estrogenic activities. In previous studies, ERβ activation in MB and CNS-PNET tumor cells was found to stimulate cytoprotective mechanisms which decreased caspases 3 activity, and loss of ERβ activity inhibited MB tumor growth and increased apoptosis *in vivo* [27]. The results of the current study lend additional evidence that ERβ initiated mechanisms promote MB and CNS-PNET survival by demonstrating that the ERβ selective agonist DPN protected D283 Med cells from cisplatin induced cytotoxicity, and that inhibition of ERβ by PHTPP blocked the protective effect of E2. Along with demonstrating mechanistic involvement of ERβ, the ability of the anti-estrogen chemotherapeutics fulvestrant and tamoxifen to each block the protective actions of estrogen in MB and CNS-PNET support previous findings which demonstrated that antiestrogen chemotherapeutics block the growth of MB tumors in vivo [26, 27] and that tamoxifen can sensitize MB cells to the cytotoxic effects of the topoisomerase inhibitor etoposide [43].

Specific treatments for MB and CNS-PNETS is constantly evolving. Depending on specific risk stratification, the current standard of care often includes maximal surgical resection that allows preservation of neurological function, postoperative radiation therapy, followed by chemotherapy employing a combination of the DNA crosslinking agent cisplatin, a DNA alkylating agents such as lomustine, and vincristine, a microtubule inhibitor [21, 44]. Each of these drugs works in different ways to stop the growth of tumor cells, either by killing the cells, or by stopping them from dividing. As for other cancers, treatment for MB and CNS-PNET has leveraged the fundamental understanding that cancer patients given multimodal treatments which include some combination of surgical tumor resection, radiation, plus a single or multiple chemotherapy agents, have improved short and long term outcomes with the combine effect of multiple cytotoxic treatments arising because each targets different processes involved in tumor cell survival [45, 46]. Targeted cancer therapies based on molecular markers such as endocrine therapy for prostate cancer or inhibiting ER activity in ER-positive breast cancer also benefits from a multimodal treatment approach that can includes endocrine based therapies, along with chemotherapy involving single or multiple cytotoxic agents [19, 20, 47–51]. Because E2 increases MB tumor survival through a general cytoprotective mechanism by increasing IGF-signaling [27], we hypothesized that its cytoprotective effects would decrease the cytotoxic effects of chemotherapeutic agents used for MB treatment independent of their mechanism of action. The presented studies, focused primarily on the most commonly used MB chemotherapy drug cisplatin (a DNA crosslinking agent), found that estrogen and soy-derived phytoestrogens were modestly cytoprotective, typically causing about a 2 fold increase in viability. The cytotoxic effects of both the alkylating agent lomustine and the microtubule inhibitor vincristine were also decreased by E2 in D283 Med MB cells demonstrating that the estrogen-induced cytoprotective mechanisms were in fact independent of the mechanism by which these chemotherapeutic agents act to initiate MB cell death.

We and others have found that Daoy cells express ERβ, but the pattern of ERβ protein isoform expression is distinctive from other MB and CNS-PNET cells in which E2 has cytoprotective activities [26, 52]. We also previously found that estrogen stimulated the migration of Daoy cells by an ERβ-dependent mechanism that was identical to that observed in other MB and CNS-PNET cells [26]. It was found here however, that unlike other MB cells, E2 alone did not increase viability of Daoy cells, instead the presence of E2 increased sensitivity of these cells to cisplatin cytotoxicity. It was also observed that inhibition of estrogen activity with 10 nM fulvestrant (a concentration that is 10-fold more than required to fully inhibit estrogen dependent growth of MCF-7 breast cancer cells [53]) blocked estrogen mediated sensitization of Daoy cells to cisplatin. At this fulvestrant concentration the classical nuclear receptor transactional activities of the ERs are inhibited, suggesting that blockage of ERβ activity is responsible for the observed chemoresistance to cisplatin, fulvestrant however, also acts as a full agonist of rapid ERβ signaling in cerebellar granule cell precursors [54]. The impact of fulvestrant agonist activity on rapid estrogen signaling in MB remains to be clearly defined.

Urbanska and colleagues, while not investigating the growth promoting actions of estrogen, previously reported that ERβ could interact with nuclear IRS1 to inhibit Rad51 mediated DNA repair mechanisms in Daoy cells, findings that suggested estrogen’s ability to increase Daoy cell sensitivity to cisplatin might involve an ERβ/IRS1 mediated decrease in Rad51 homologous recombination DNA repair mechanisms [52]. Their additional results from experiments using a much higher 10 μM concentration of fulvestrant (IC_50_ = 0.29 nM) found it caused resistance of Daoy and D384 cells to the cytotoxic actions of cisplatin, effects that were not significant in the D283 Med cells [34]. The studies presented here, using lower concentrations of fulvestrant, failed to observe increased sensitivity of Daoy cells to cisplatin. In light of the higher concentrations of fulvestrant used for those previous experiments, it is possible that the observed protective effects of fulvestrant were not specific and involved activities other than inhibition of ERβ. Another possible explanation for decreased cytotoxicity could be that DMSO, if used as a vehicle, was decreasing the efficacy of cisplatin. The ability of DMSO to greatly decrease the cytotoxic activity of cisplatin and other platinum chemotherapeutic drugs has previously been characterized in detail [35]. It is also valuable to consider the contracting results observed here, and in other previous studies, in view of the fact that current molecular profiling and cytogenetic data supports the conclusion that the Daoy cell line, while most closely resembling the SHH molecular subgroup, is markedly different from all primary MB tumor subgroups and distinctive from MB cell lines like D283 Med that retain the hallmarks of MYC amplification and i17q which are associated with poor outcomes [55, 56]. It is especially notable that the hypertetraploid karyotype of the Daoy cells does not resemble MB karyotypes, and the presence of two X chromosomes and a lack of a Y chromosome, is inconsistent with the sex of the male patient from which the original tumor biopsy was isolated [42]. In spite of these caveats, it is possible that the observed differences in the impact of E2 on Daoy and D283 Med MB cells may actually reflect the well-known heterogeneity of MB. It is likely that different molecular subtypes of MB and even different populations of cells in a single patient’s tumor may differently respond to ER agonists and antagonists. This raises the interesting possibility that the responses to estrogen in MB are heterogeneous and that some populations of cells are differentially responsive to estrogen's effects.

## Conclusions

The presented results demonstrate that the cytoprotective effects of E2, which can be cell line dependent, are clearly chemoprotective in some MB and CNS-PNET cell lines. The results of additional experiments also demonstrated that like E2, low and physiological levels [41] of the soy-derived phytoestrogens genistein, daidzein, and equol can decrease caspase activity in D283 Med MB cells resulting in an estrogen-like inhibition of the cytototoxic actions of cisplatin. The finding that soy phytoestrogens also decrease sensitivity to the cytotoxic actions of cisplatin suggest that attention to decreasing exposures to environmental estrogens that include not only phytochemicals but also xenobiotic endocrine disruptors may benefit MB patients undergoing cytotoxic chemotherapy.

## Declarations

### Ethics approval and consent to participate

N/A

### Consent for publication

N/A

### Availability of data and material

All data generated or analyzed during this study are included in this published article

### Authors’ contributions

SB analyzed and interpreted data, performed cell viability experiments and wrote the manuscript; CB performed and analyzed cell viability and apoptosis experiments; CC performed cell viability assays, assisted in data collection and analysis and contributed early drafts of the manuscript; MK performed and analyzed cell growth studies, GM, FS and KW performed and analyzed cell viability studies.

## Acknowledgements

As part of the Department of Pharmacology and Cell Biophysics, Summer Undergraduate Research Program, Katie Wray was a Dalton Zannoni Summer Undergraduate Research Fellow of the American Society of Pharmacology and Experimental Therapeutics. We are grateful for the assistance of Drs. Robert Rapaport and JoEl Schultz and Nancy Thyberg in support of graduate and undergraduate training.

## List of abbreviations

(MTS): 3-(4,5-dimethylthiazol-2-yl)-5-(3-carboxymethoxyphenyl)-2-(4-sulfophenyl)-2H-tetrazolium
(MTT): 3-(4,5-dimethylthiazol-2-yl)-2,5-diphenyltetrazolium bromide
(E2): 17β-estradiol
(CNS-PNET): central nervous system primitive neuroectodermal tumors
(CSS): charcoal stripped FBS
(DMSO): Dimethylsulfoxide
(EBSS): Earle’s Balanced Salt Solution
(FBS): fetal bovine serum
(MB): medulloblastoma
(MBEN): medulloblastoma with excessive nodularity
(MEM): minimum essential media
(pNA): p-nitroaniline
(PNETs): primitive neuroectodermal tumors
(SHH): sonic hedgehog
(WNT): wingless

